# Pangenome-based dynamic trajectories of intracellular gene transfers in Poaceae unveil a high rate of unceasing integration and selective retention in Triticeae

**DOI:** 10.1101/2022.10.11.511703

**Authors:** Yongming Chen, Yiwen Guo, Xiaoming Xie, Zihao Wang, Lingfeng Miao, Zhengzhao Yang, Yuannian Jiao, Chaojie Xie, Jie Liu, Zhaorong Hu, Mingming Xin, Yingyin Yao, Zhongfu Ni, Qixin Sun, Huiru Peng, Weilong Guo

## Abstract

Intracellular gene transfers (IGTs) between the nucleus and organelles, including plastids and mitochondria, constantly reshapes the nuclear genome during evolution. Despite the substantial contribution of IGTs to genome variation, the dynamic trajectories of IGTs at the pangenomic level remain elusive. Here, we propose a novel approach, IGTminer, to map the evolutionary trajectories of IGTs by collinearity and gene reannotation across multiple genome assemblies. IGTminer was applied to create a nuclear organelle gene (NOG) map across 67 genomes covering 15 Poaceae species, including important crops, revealing the polymorphisms and trajectory dynamics of NOGs. The NOGs produced were verified by experimental evidence and resequencing datasets. We found that most of the NOGs were recently transferred and lineage specific, and that Triticeae species tended to have more NOGs than other Poaceae species. Wheat had a higher retention rate of NOGs than maize and rice, and the retained NOGs were likely involved in the photosynthesis and translation pathways. Large numbers of NOG clusters were aggregated in hexaploid wheat during two rounds of polyploidization and contributed to the genetic diversities among modern wheat varieties. Finally, we proposed a radiocarbon-like model illustrating the transfer and elimination dynamics of NOGs, highlighting the unceasing integration and selective retention of NOGs over evolutionary time. In addition, we implemented an interactive webserver for NOG exploration in Poaceae. In summary, this study provides new resources and clues for the roles of IGTs in shaping inter- and intraspecies genome variation and driving plant genome evolution.

## INTRODUCTION

In plant cells, the nucleus and two organelles, chloroplasts (plastids) and mitochondria, originating from prokaryotic endosymbiotic ancestors, together make up the genetic material (1). It has been accepted that the genetic material has been transferred among these three genetic compartments (1-4). Organelle-to-nucleus DNA transfer is considered to have the highest frequency among all types of intracellular DNA transfers, ubiquitously and continually reshaping nuclear genomes over evolutionary time (5-8). The organelle genome contains important functional genes covering ribosomal proteins, ATP synthase, and photosynthesis pathway (3, 9); thus, intracellular gene transfer (IGT), also known as endosymbiotic gene transfer (EGT), provides potential functional copies of these genes for plants (8). Organelle-to-nucleus fragment integration via nonhomologous end-joining during the double-strand break repair process might be a common mechanism conserved in eukaryotic organisms (5, 10). Once integrated into the nuclear genome, organelle-derived DNA can experience different fates, including elimination, rearrangement, fragmentation, and mutation (10-12). Insertions of cytoplasmic DNA generally occur in clusters and can be large in size (13, 14). They can also be structurally complex, with multiple fragments from different regions often merging into the same nuclear genome location (13, 15). Nucleus organellar DNA transfer content is positively correlated with the nuclear genome size (10) and the number of mitochondria or plastids in the cell (16, 17). To date, many studies of organelle-to-nucleus transfer of genetic material in plants have been reported (10, 18-25), most of which focused on the sequences of DNA fragments, which are generally referred to as nuclear plastid DNAs (NUPTs) and nuclear mitochondrial DNAs (NUMTs), and were irrespective of fragment length. NUPTs may have an important contribution to shaping the sex chromosome centromere of *Asparagus officinalis* (26). NUPTs and NUMTs may contribute to genetic diversity and environmental adaptation in rice accessions (24). A recent study found that a 1.64 Mb insertion containing transgenic insertions in SunUp papaya, consisted of 52 NUPTs, 21 NUMTs, and 1 nuclear genomic fragment (27).

Some studies have performed organelle genome transformation and editing to improve the yield, nutritional quality, and environmental adaptation of plants (28-30). The nuclear origin supplementation of the D1 protein, encoded by the chloroplast gene *psbA*, increases the photosynthetic efficiency of plants (28, 31). Simultaneous chloroplast transformation of Rubisco large and small subunits can modify plant photosynthesis and growth (32). In addition, the increased translation efficiency of photosystem II contributes to the adaptation of *Arabidopsis thaliana* to high light environments (33). Horizontal gene transfer (HGT) (8, 34) and IGT can introduce more functional gene copies and increase genomic diversity, which may provide organisms with functions or phenotypes to adapt to diverse environments. A recent study demonstrated that the reductive evolution of organelle genomes and IGTs are likely to be associated with reducing cellular cost and improving energy efficiency (35).

Poaceae contains many economically important cereal crops, such as bread wheat (*Triticum aestivum*), durum wheat (*Triticum turgidum*), barley (*Hordeum vulgare*), rye (*Secale cereale*), oat (*Avena strigosa*), rice (*Oryza sativa*), and maize (*Zea mays*). Some species have large and complex genomes. For example, bread wheat is an allopolyploid (BBAADD) with an extreme genome size (16 Gbp per haploid genome) (36). With the development of genome assembly algorithms and strategies, the number of available well-assembled genomes in Poaceae has increased rapidly in recent years (37). Pangenome projects for wheat (38) and barley (39) and rice gap-free genomes (40) have been completed. However, the dynamic trajectories of IGTs at the pangenomic level remain largely unknown. These resources have pushed genome research for Poaceae into the pangenome era and can provide great opportunities for obtaining new insights into the role of IGTs in shaping genome content variation.

Collinearity across genomes can provide insights into evolutionary trajectories of genes that are connected to presence/absence variations (41, 42). A homology inference method based on collinearity was developed for Triticeae and showed that α-gliadin genes are typically tandemly duplicated with an expansion of clusters during the evolution of bread wheat (41). The contribution of tandem duplication and whole-gene duplication to gene family expansion in the early evolutionary history of the CIPK gene family has been reported (43). These studies demonstrate the power of collinearity in tracking the evolutionary history of genes across multiple genomes. In addition, the gene annotation quality of assemblies is of key importance for the phylogenomic study of genes (44, 45). Current gene annotations in the nucleus genome are mainly automatically produced by universal pipelines, such as MAKER-P (46). However, these traditional annotation methods may miss genes with short coding sequences and variable start codons. Some gene annotation tools for organelle genomes, such as DOGMA (47), MITOS (48), and PGA (49), were specifically designed to perform plastid or mitochondrion genome annotation but are not yet suitable for the nuclear genome. A new strategy for exploring and connecting IGTs is urgently needed to address pangenomic research challenges.

Here, we present the first study of the trajectory dynamics of IGTs at the pangenomic level in plants. A novel method, IGTminer, were developed to detect and connect IGTs across multiple assemblies by gene reannotation and collinearity. We constructed a Poaceae nuclear organelle gene (NOG) map using 67 genomes covering 15 Poaceae species. We found that the Triticeae species tended to have more NOGs than other Poaceae species and that wheat had a higher rate of NOG retention than maize and rice. We also investigated the evolutionary history of NOGs during polyploidization and the diversity among wheat populations. Finally, we proposed a radiocarbon-like model illustrating the dynamics of NOGs and implemented an online platform for NOG exploration in Poaceae. In general, our work provides new insights into the role of IGTs in shaping inter- and intraspecies genome variation in plants.

## RESULTS

### IGTminer incorporates gene reannotation and collinearity information to explore the dynamical trajectories of IGTs across assemblies

To characterize the evolutionary trajectories of IGTs (**Figure 1A**) in Poaceae, we first investigated the genomic distribution of NUPTs and NUMTs in important cereal crops, including wheat (50), barley (39), rice (51), and maize (52) (**Figure 1B**). The results showed that the average total length per genome is 1.4 Mb (range 0.9–2.4 Mb) for NUPTs and 3.8 Mb (range 0.8–7.1 Mb) for NUMTs. However, on average, only 6.2% of NUPTs and 1.9% of NUMTs contain gene sequences, and most NUMT/NUPT sequences are generally short. In addition, we performed a phylogenetic screen in these genomes and found that this method produced similar IGTs to the single-genome screen (mean Jaccard similarity = 0.963). The above findings are consistent with previous studies that most of the NUMT/NUPT fragments acquired from organellar genomes would be eliminated and mutated gradually over evolutionary time (7, 10). We only focused on fragments containing nuclear plastid genes (NPGs) or nuclear mitochondrial genes (NMGs) in Poaceae.

**Figure 1.**
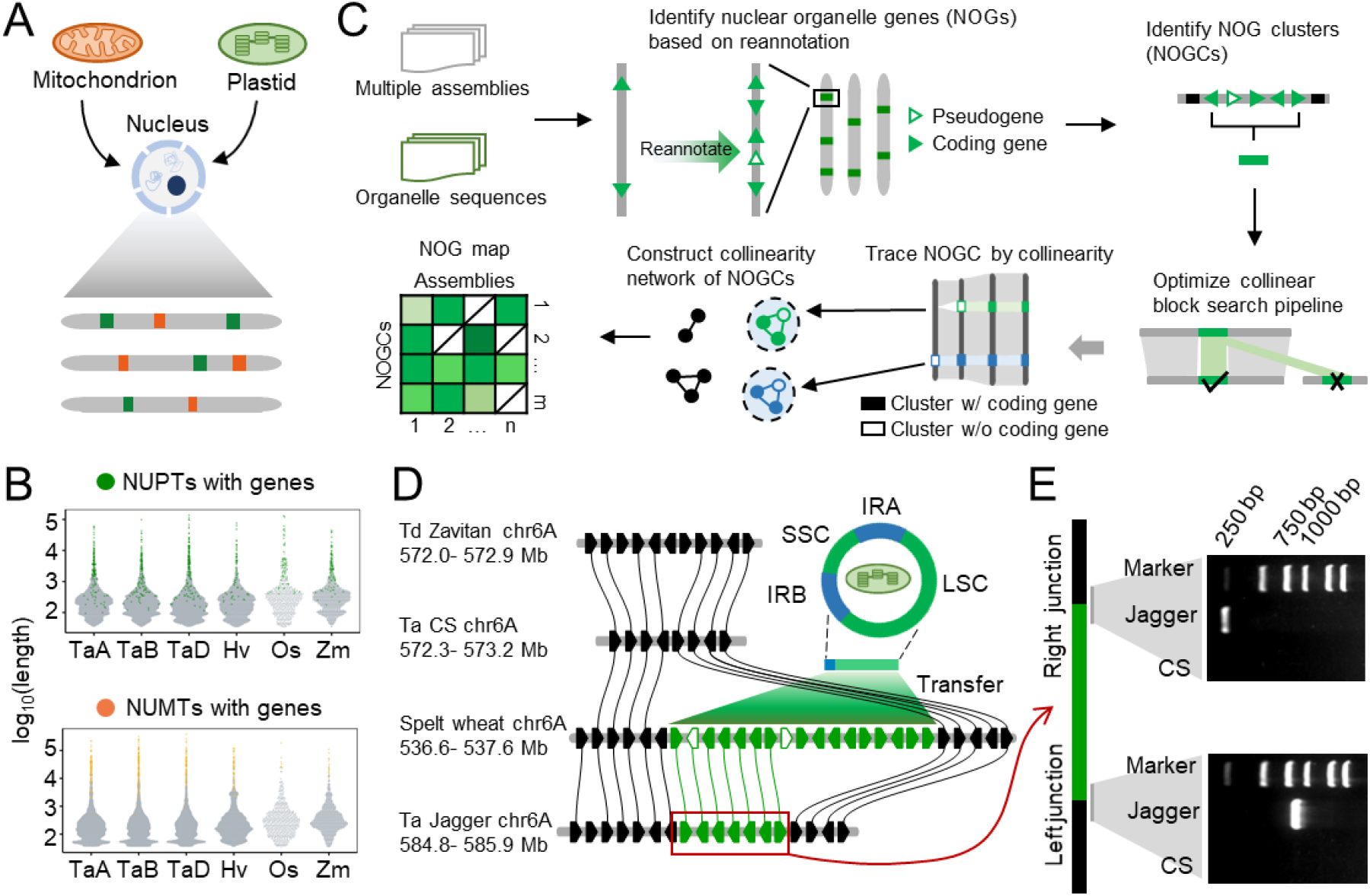
The IGTminer pipeline for connecting NOGs across assemblies and recording their dynamic trajectories. **(A)**A schematic diagram of organelle-to-nucleus gene transfer. **(B)**The length distribution of NUPTs and NUMTs in the genomes of the main cereal crops. Y-axis, log2-transformed lengths in base pairs. Colored dots represent NUPTs and NUMTs containing annotated genes. Gray dots represent NUPTs and NMPTs without annotated genes. TaA, TaB, and TaD represent the A, B, and D subgenomes of bread wheat, respectively. Hv, *Hordeum vulgare*. Os, *Oryza sativa*. Zm, *Zea mays*. **(C)**Design of the IGTminer pipeline for uncovering the evolutionary trajectories of the IGT (NOG) using gene reannotation and collinearity information. First, raw NOG annotations in the reference genome are removed and reannotated. The reannotated genes are then clustered into a single transfer event. The collinearity block search pipeline is optimized and used to search collinear NOGs. Only NOGC networks containing at least one coding gene are used. Finally, a NOG map across multiple genomes recording the dynamics of NOGs is generated. **(D)**A typical example showing polymorphism and collinearity of NPGs between four genomes. Genomes of two bread wheat accessions (cv. Chinese Spring and Jagger), a wild emmer wheat accession (Zavitan), and a spelt wheat accession are presented. An unfilled pentagon represents a pseudogene, and a filled pentagon represents an annotated coding gene. Green pentagons, NPGs. Black pentagons are nucleus-oriented genes. Lines represent orthology relationships. The direction of the spelt wheat track was reversed manually. LSC, large single-copy region. SSC, small single-copy region. IRA and IRB, a pair of inverted repeats. **(E)**Confirmation of the presence of the NPGC in the nuclear genome with PCR amplification experiments. PCR products span plastid DNA-nucleus junction sites on both sides of the NPGC. Black block, nuclear DNA. Green block, NPGC DNA.

We designed a robust tool, IGTminer, by incorporating gene reannotation and collinearity information to unveil the evolutionary trajectories of IGTs across multiple genome assemblies (**Figure 1C**). IGTminer includes the following main steps. First, NOGs in reference genomes are reannotated. Many genomes are independently annotated by diverse pipelines, which results in low comparability across multiple assemblies. To improve the quality of NOG annotation, we reannotated all NOGs in the reference genome using the homology-based annotation method. The resulting gene models are further classified into coding genes and pseudogenes. Second, NOG clusters (NOGCs) are identified based on neighboring genome locations. Some NOGs commonly exist in single clusters because they are derived from a single transfer event. Multiple NOGs with short intervals are clustered into several single nuclear integration events, which can be classified into the NPG cluster (NPGC) and the NMG cluster (NMGC). Third, we improved the pairwise collinear block search pipeline for identifying collinear NOGCs. A polyploid assembly is decomposed into multiple diploid subassemblies to meet the need for phylogenomic analysis. To avoid issues with the inaccuracy of collinear block search caused by high-similarity NOGCs, we substitute NOGCs into single sites to improve the search power of the collinear NOGCs. Fourth, the Markov cluster algorithm (MCL) (53) is applied to construct the collinearity networks of NOGCs, where each node represents an NOGC and each edge represents the collinearity between a pair of NOGCs from different genomes. The NOGC networks without any coding genes are filtered to make the NOG dataset more accurate and less redundant. Finally, IGTminer takes multiple assemblies as input and produces a NOG map, recording the trajectory dynamics of NOGs at the pangenomic level.

### Evaluation of IGTminer and experimental validation of identified NOGs

We evaluated the performance of IGTminer by taking genome assemblies as input with whole-genome resequencing (WGS) data. The WGS sets with an average genome coverage of ∼6× from eleven bread wheat accessions were acquired from previous studies (38, 41). An NOGC site identified by IGTminer is real only when the WGS reads spanning the organelle DNA-nucleus junction sites are detected (**Figure S1**). The results show that an average of 99.0% of NPGCs and 99.4% of NMGCs identified by IGTminer using genome assemblies could be recalled by WGS data. In addition, all the sequences of identified NOGs in wheat, barley, rice, and maize aligned with best hits of plastid or mitochondrial sequences in the UniProt Swiss-Prot protein database, indicating that the NOGs identified by IGTminer are of plastid or mitochondrial origin.

Experiments were also conducted to validate several NOGCs identified by IGTminer. A plastid integration on wheat cv. Jagger was found to be well collinear with the plastid genome (**Figure 1D**). By tracing this NPGC across the hexaploid wheat and wild emmer wheat genomes (54), only one collinear region in the spelt wheat genome was found. PCR amplification experiments were then conducted. The results showed that all the amplified fragments covered plastid DNA-nuclear junction sites, which verified the presence of NPGC (**Figure 1E**). In addition, an NMGC in the CS genome had no collinear region in other genomes (**Figure S2A**), which was also confirmed by PCR amplification experiments (**Figure S2B**). In summary, IGTminer is an ideal approach for NOG map construction using a panel of assemblies.

### Landscapes and diversities of NOGs in Poaceae

To generate a broad-scale NOG profile, we collected a total of 67 representative and high-quality genome assemblies (**Table S1**) from 15 Poaceae species covering important cereal crops, such as wheat (41, 50, 55), rye (56, 57), barley (39), oat (58), rice (40, 51, 59, 60), sorghum (61), and maize (25, 52, 62-66). After decomposing the assemblies of polyploid species, we obtained a total of 93 diploid assemblies. The IGTminer pipeline was applied to produce a Poaceae NOG map recording the dynamic trajectory of 1,302 NPGCs (**Figure 2A**) and 431 NMGCs (**Figure S3A**). We found that the counts of genes (i.e., NOGs) and transfer events (i.e., NOGCs) differed across diploid assemblies in Poaceae. In detail, NPGs ranged from approximately 28 to 403 (14-fold difference) (**Figure 2B**) and NMGs ranged from approximately 22 to 133 (6-fold difference) (**Figure S3B**). NPGCs ranged from approximately 13 to 93 (7-fold difference) (**Figure 2C**) and NMGCs ranged from approximately 6 to 60 (7-fold difference) (**Figure S3C**). The counts of NPGs were always larger than those of NMGs, which is likely associated with the higher chloroplast copy number per cell. Of all the Triticeae species, barley and hexaploid wheat had the lowest and highest counts of NOGs, respectively. We next investigated the divergence of NOGs across three wheat lineages and found that the wheat D lineage (370 genes on average) had more NPGs and NPGCs than the A lineage (172 genes on average) and B lineage (166 genes on average) (**Figure S4A and S4B**). Unlike plastid-to-nucleus gene transfer, the wheat B lineage (133 genes on average) had more NMGs and NMGCs than the A lineage (107 genes on average) and D lineage (115 genes on average) (**Figure S4C and 4D**). Triticeae species have a higher rate of plastid gene transfers (**Figure 2D**) and mitochondrion gene transfers (**Figure S3D**) than others in the Poaceae.

**Figure 2.**
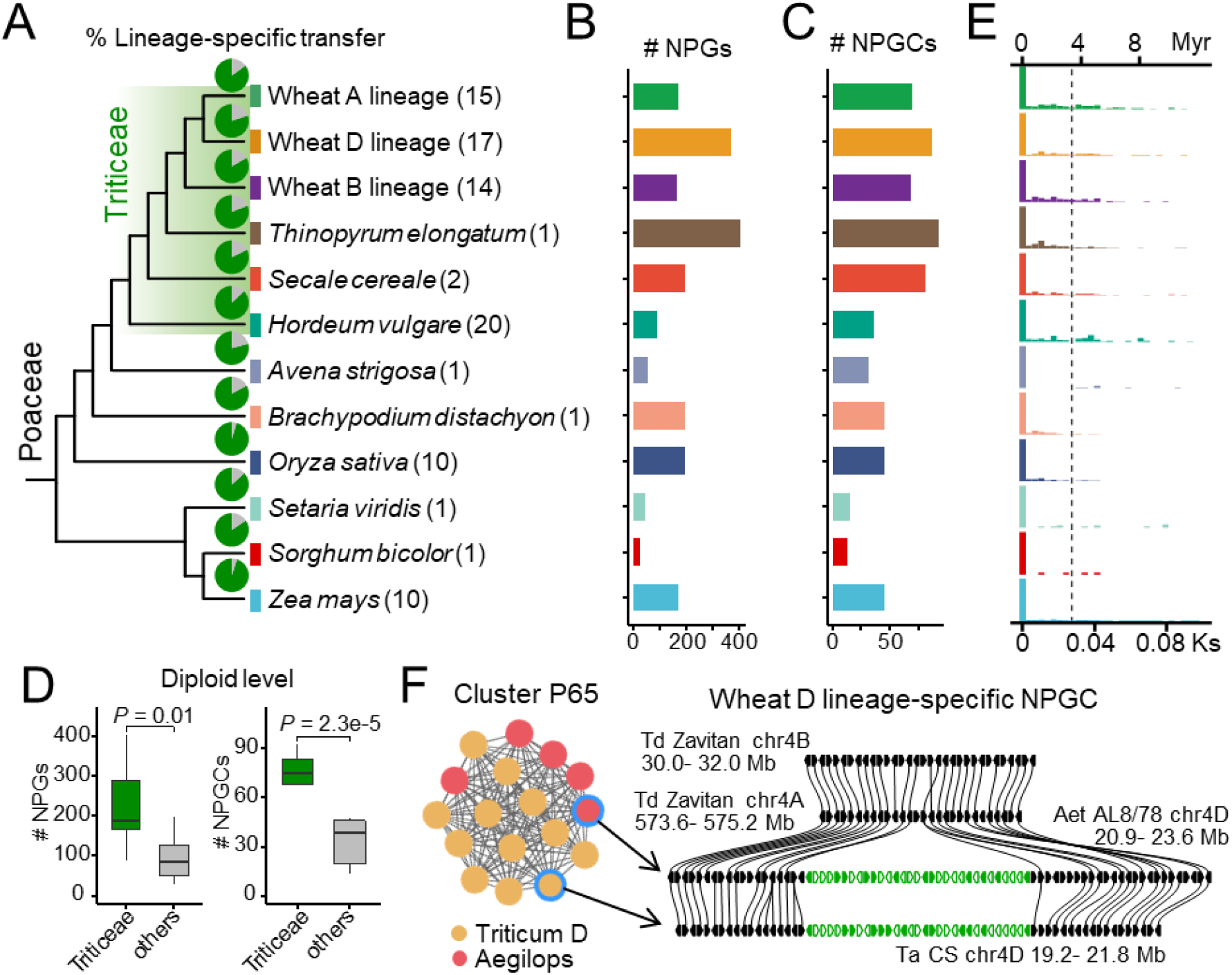
Overview and diversity of NPGs in Poaceae. **(A)**Phylogenetic tree of the 12 Poaceae species/lineages. The numbers in parentheses show the counts of the genomes used in each lineage. The left tree is supported by previous studies (70, 81, 92) and the TimeTree Website (http://timetree.org/). A total of 93 diploid assemblies were used. The tribe Triticeae was highlighted. The green pies show the proportion of lineage-specific NPGCs in each lineage. **(B and C)** NPG (B) and NPGC (C) average counts in each lineage. **(D)** Comparison of counts of plastid-to-nucleus gene transfers between Triticeae and others in Poaceae. The Wilcoxon test was performed. **(E)** The estimated transfer times of NPGs, as investigated by the Ks method. The dashed line indicates the differentiation time of the wheat A and D lineages. The tribe Triticeae evolved approximately 25 million years ago in the Poaceae (93). **(F)** An example shows the wheat D lineage-specific NPGC network. Cluster P65 is highly conserved in the wheat D lineage. In the left network, each node represents an NPGC, and edges indicate collinearity relationships. Red nodes represent NPGCs from diploid Aegilops genomes, and orange nodes represent NPGCs from the D subgenome of hexaploid wheat. The right microcollinearity plot shows the polymorphism and collinearity of NPGCs among the four genomes. An unfilled pentagon represents a pseudogene, and a filled pentagon represents an annotated coding gene. Green pentagons, NPGs. Black pentagons are nucleus-oriented genes. Lines represent orthology relationships. The direction of the Zavitan track was reversed manually. The genomes shown in the microcolinearity plot are highlighted with blue rings in the network.

### Most NOGs have recently emerged and are lineage-specific in Poaceae

To further estimate the emergence time of NOGs, we first utilized the widely used Ks (number of substitutions per synonymous site) method (67, 68). The divergence times between species were consistent with those reported in previous studies (69, 70) (**Figure S5**). Furthermore, the transfer time analysis of NPGs (**Figure 2E**) and NMGs (**Figure S3E**) showed that most NPGs (83.1% on average) and NMGs (86.3% on average) in Poaceae occurred after the differentiation of the wheat A, B, and D lineages, thus species of Poaceae have recently abundant organelle-to-nucleus gene transfers. Evolutionary conservation can provide many clues concerning the emergence time of a gene in an organism (41, 43). Thus, we investigated the collinearity of NOGCs across Poaceae revealed by IGTminer. We found that approximately 85.3% of NPGCs (**Figure 2A**) and 95.0% of NMGCs (**Figure S3A**) are lineage-specific across Poaceae, indicating that organelle-to-nucleus gene transfers are poorly conserved, which can also be supported by the short transfer times and low Ks values of NOGs (**Figure 2E and S3E**). For instance, the well-collinear NPGC (**Figure 2F**) and NMGC (**Figure S6**) were only located in the wheat D lineage and B lineage, respectively.

### Integration preference of NOGs in Triticeae

To obtain a new understanding of the transfer preference of NOGs at the pangenomic level, we first profiled the chromosome positions of 409 NPGCs and 285 NMGCs in the wheat pangenome **(Figure 3A)**, and found that NOGs were distributed on all wheat chromosomes. In addition, by investigating the CS genome alone, we found that NPGCs and NMGCs were often located in adjacent chromosome positions (**Figure 3B**). Previous studies have reported that large NUPTs tend to be distributed in the centromeric and pericentromeric regions in *Asparagus officinalis* (26) and rice (71). In this study, NOGCs in wheat had a higher frequency of location at the end of the long arm and short arm (**Figure 3C**), which may be divergent from *As. officinalis* and rice.

**Figure 3.**
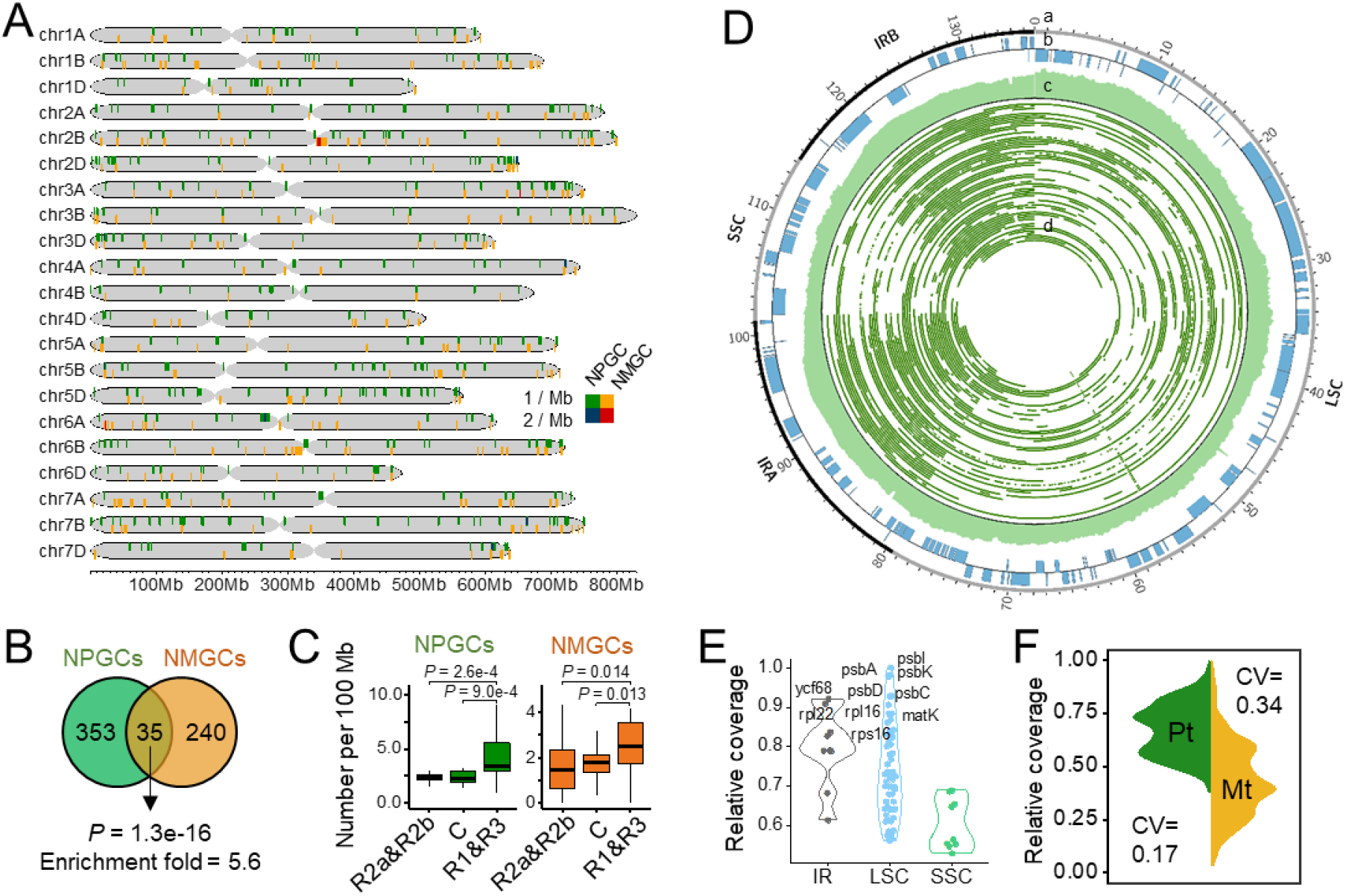
Distribution and transfer preference of NOGs in Triticeae. (**A**) Positions of all NOGCs in the bread wheat nucleus genome. Green and blue, NPGC. Orange and red, NMGC. Green and orange, one event per Mb. Blue and red, two events per Mb. NOGCs in eleven hexaploid wheat genomes were anchored to the Chinese Spring reference genome with collinearity. (**B**) Colocalization of NPGCs and NMGCs in the Chinese Spring wheat genome. The adjacent NOGCs were determined by an interval of 1 Mb. A hypergeometric test was performed. (**C**) Density of NOGCs on different chromosome zones, as defined by a previous study (50, 94). The Wilcoxon test was performed. R1 and R3 represent distal regions of the chromosomal long arm and short arm, respectively. C represents the centromeric/pericentromeric region. R2a and R2b represent the rest. (**D**) Circos plot shows the distributions of original plastid sequences of NPGCs. Tracks show chromosomal position (a), gene position (b), density of mapped NPGC sequences (c), and mapping position of NPGC sequences (d). In track a, the numbers represent the length in kilobases. In track b, Medial, reverse strand; Lateral, forward strand. In track c, the mapping positions of NPGC sequences are normalized with repeat sequence abundance. (**E**) NPGC coverages on original plastid genes in three regions, LSC (large single-copy), SSC (small single-copy), and IR (inverted repeat) regions. Each point represents one gene. The symbol names of the top ten genes with the highest coverage are noted. For easier comparison of plastid and mitochondrion genomes, the coverage of each site was normalized by dividing by the coverage of the site with the highest transfer frequency. (**F**) Comparison of coverage of integration sequences in different bins of the original plastid (pt) and mitochondrial genomes (mt). For NPGCs and NMGCs, we normalized the coverage of each site, referring to the method in Figure 3E. Coefficient of variation, sd of coverage/mean of coverage.

Motif enrichment analysis was performed to investigate the sequence features of the transfer sites. A previous study of rice showed that organellar sequences might prefer to insert into AT/TA repeat-enriched regions in the nucleus (24). We also obtained AT/TA repeat enrichment in the flanking region of NOGCs in the rice genome using the same pipeline, while there was no enrichment of repeats in wheat genomes (**Figure S7**). We further counted the AT content of flanking regions of NOGCs, and found significantly higher levels for NPGCs (**Figure S8A**) and NMGCs (**Figure S8B**).

The preference analysis of the original organelle genes of the NOGs (**Table S4; see methods**) showed that the top ten plastid genes with the highest transfer frequency were mainly located in the LSC region (**Figure 3D and 3E**), and most of those genes are involved in photosynthesis and translation pathways. The SSC region has the lowest transfer frequency, indicating the existence of imbalanced transfer from the plastid to the nucleus. We also analyzed the original genes of NMGs (**Figure S9A**) and found that the top ten mitochondrial genes with the highest transfer were mainly classified as ribosomal proteins, NADH dehydrogenase subunits, and ATP synthase subunits (**Figure S9B**). It was interesting that the coefficient of variation (0.34) of NMGCs was higher than that of NPGCs (0.17) in terms of the transfer frequency (**Figure 3F**), indicating that the transfer sequence of plastids tends to be more complete. The original sequences of NPGCs are generally longer than those of NMGCs (**Figure S10**). These observations highlight the imbalanced transfer preference of the organelle sequence.

### Large numbers of NOGs were retained in hexaploid wheat during polyploidizations

Bread wheat has undergone two rounds of polyploidization (72), which increases genomic complexity. However, the dynamics of NOGs during this process remain unknown. We thus evaluated the evolutionary conservation of NPGCs across bread wheat (three subgenomes), *T. turgidum, T. urartu*, and *Ae. tauschii* (**Figure S11A and S11C**). The results show that the origin of half or more NPGCs can be traced to ancestor genomes (**Figure 4A**). For instance, the NPGC clusters P110 and P155 are representative transfers that occurred before the formation of the *T. urartu* and *T. turgidum* A subgenomes, respectively (**Figure 4A**). The NPGC clusters P108 and P248 are representative transfer events before the formation of the *Ae. tauschii* and *T. aestivum* D subgenomes, respectively (**Figure S12**). The NPGC clusters P168 and P347 are representative transfer events during wheat B genome evolution (**Figure S13**). In addition, the presence of *T. urartu*- and *Ae. tauschii*-specific NPGCs suggests that a portion of NPGCs in the diploid ancestor genomes of wheat did not pass to bread wheat.

**Figure 4.**
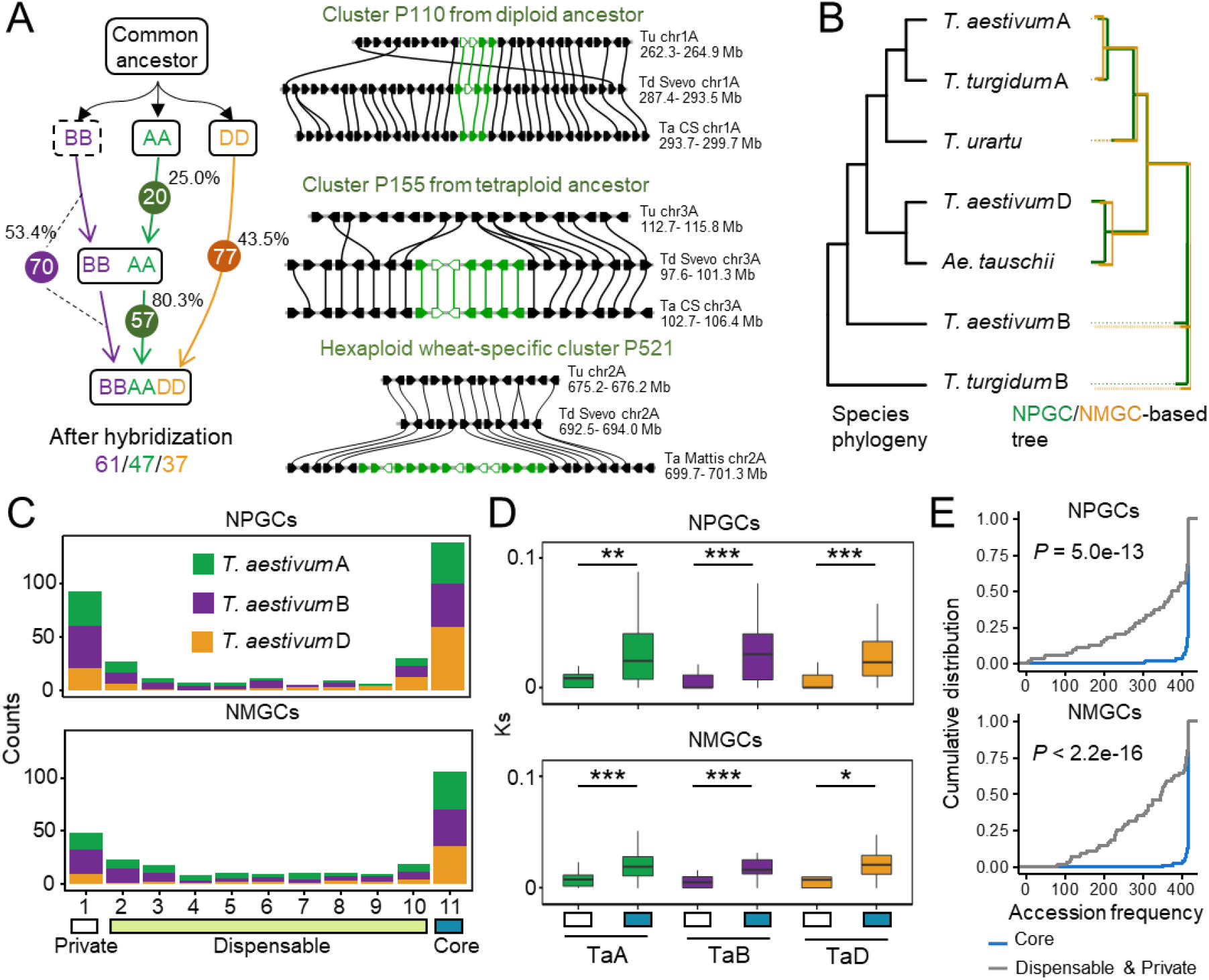
Dynamic trajectories of NOGs during wheat evolution. (**A**) Overview of the retained NPGCs in wheat during two rounds of polyploidization. Numbers in circles indicate the number of retained NPGCs. The percentages next to the numbers represent the retention ratio of NPGCs during polyploidization. Two examples show representative transfer events during the evolution of the wheat A genome. Microcollinearity plots show polymorphism and collinearity of NPGCs among multiple wheat A genomes. An unfilled pentagon represents a pseudogene, and a filled pentagon represents a coding gene. Green pentagon, NPGs. Black pentagons, nucleus-oriented genes. Lines represent orthology relationships. (**B**) NOGCs-based phylogenetic tree. The left tree is supported by previous studies (70, 81, 92) and the TimeTree Website (http://timetree.org/). The right tree indicates the tree constructed using the presence/absence variants (PAV) matrix of NPGCs (green) and NMGCs (orange) across multiple genomes. (**C**) Counts of the NPGCs and NMGCs shared by eleven wheat genomes. The classification of core, dispensable, and private sets was performed according to a previous study (60). (**D**) Ks distributions of NOGs in private and core sets. The Wilcoxon test was performed. *, P < 0.05. **, P < 0.01. ***, P < 0.001. (**E**) Cumulative distributions show the occurrence frequencies of NOGC across wheat accessions. The Wilcoxon test was performed. Resequencing datasets of 418 bread wheat accessions were used.

Based on the above findings, we expected to re-infer the evolutionary history of bread wheat with only NPGCs. With the presence/absence variants (PAV) matrix of NPGCs across bread wheat and its donor genomes, we reconstructed the NPGC-based tree (**Figure 4B**), which was consistent with the polyploidization history of bread wheat (73, 74). We also investigated the evolutionary conservation of NMGCs during polyploidization (**Figure S11B and S11D**) and re-inferred the polyploidization history of bread wheat with an NMGC-based tree (**Figure 4B**).Large numbers of NOGs in the ancestors of bread wheat have been retained in bread wheat during two rounds of polyploidization.

### The pan- and core NOGC set analysis reveals the NOG diversity across the wheat population

To determine the diversity of NOGs across wheat populations, we investigated the occurrence of NOGCs in eleven bread wheat assemblies. We found that the NOGC-based tree (**Figure S14**) of ten bread wheat accessions in the 10+ Genome Project reflects well the genetic relationships among accessions (38). The NOGCs were classified into core (i.e., present in all accessions), dispensable (i.e., present in more than one but < 11 accessions), and private sets (i.e., present in one accession alone), according to the criteria that were defined in the previous study (60). The results showed that the counts of NPGCs and NMGCs in the private and core sets were much higher than those in the dispensable sets (**Figure 4C**). Ideally, the genes in the core and private sets should represent ancient (higher Ks value) and recent transfer (lower Ks value) events of organelle genes, respectively. The results revealed significant differences (*P*<0.05) between the Ks values of the private and core sets (**Figure 4D**), which validated our conjecture. We further found that most NPGCs (59.4% on average) and NMGCs (58.6% on average) were not shared across the eleven bread wheat genomes (**Figure S15**). Compared to the A (37.5% for NPGCs and 35.7% for NMGCs) and B subgenomes (31.2% for NPGCs and 31.8% for NMGCs), the D subgenomes (53.1% for NPGCs and 57.1% for NMGCs) have a higher proportion of core sets in terms of NOGCs. To further verify the dynamics of NOGCs inferred from the eleven assemblies, we acquired WGS data of 418 wheat accessions from previous studies (38, 41, 75-77), and detected NOGC polymorphisms across wheat populations (**Figure S1**). The results show the core and noncore NOGCs identified by assemblies were supported by WGS datasets (**Figure 4E**).

### Wheat has a high rate of selective NOG retention

After integration, the organelle genetic materials in the nucleus decayed during evolution, with only a small proportion being retained today (10). We found that cluster size (i.e., gene count) markedly decreased over differentiation time (i.e., Ks value) of NOGs and their original organelle genes (**Figure 5A**), which is consistent with the expectation. Furthermore, we found that the bread wheat genome has a higher proportion of NOG retention (∼23.8% of NPGs and ∼14.0% of NMGs) than other crop genomes, such as the rice (∼10.2% of NPGs and ∼1.6% of NMGs) and maize (∼17.6% of NPGs and ∼1.3% of NMGs) genomes (**Figure 5B**). In addition, we calculated the shared ratio of NOGCs across the wheat, rice, and maize assemblies based on collinearity, and found that bread wheat had a higher ratio of shared NOGCs than rice and maize (**Figure 5C**). The above results demonstrate that bread wheat retained more NOGs than others in Poaceae over evolutionary time.

**Figure 5.**
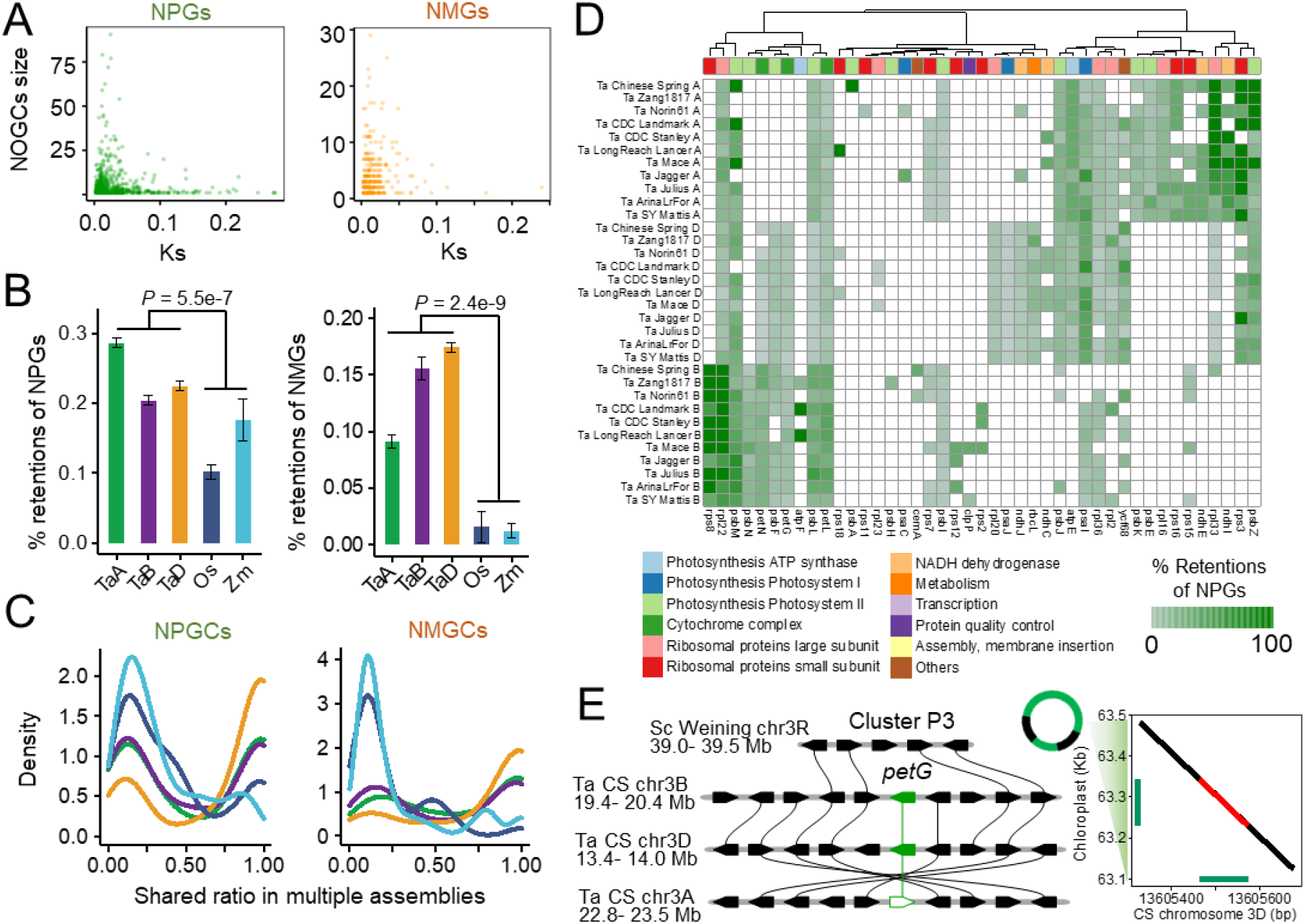
A high rate of selective retention of NOGs in wheat. (**A**) NOGC sizes decrease over time. X axis, the median of Ks for each NOGC. Y axis, the median counts of genes in each NOGC. (**B**) The proportions of retained NPGs and NMGs in the main crops such as wheat, rice, and maize. Statistics were performed on multiple assemblies (eleven wheat genomes, ten rice genomes, and ten maize genomes). The Wilcoxon test was performed. (**C**) The shared ratio of NPGCs and NMGCs revealed by wheat, rice, and maize genome assemblies. The shared ratio was calculated by dividing the counts of genomes containing the NOGCs by the total number of genomes. The colors of the lines correspond to the colors of the bars in Figure 5B. (**D**) NPG retentions in the bread wheat A, B, and D subgenomes. NPGs were classified as retained when Ks >0.04. The genes were grouped according to their biological function, as defined by a previous study (9). (**E**) The microcollinearity plot shows polymorphism and collinearity of an NPGC between four genomes. An unfilled pentagon represents a pseudogene, and a filled pentagon represents a coding gene. Green pentagon, NPG. Black pentagons, nucleus-oriented genes. Lines represent orthology relationships. Right panel, sequence alignment of the NPGC Cluster P3-located region between the 3D chromosome of the CS genome (x-axis) and the plastid genome (y-axis). Red, alignment of the *petG* gene. Green, position of the gene.

We screened the biological function of the original genes of the NOGs retained in the bread wheat genomes. First, a total of 947 retained NPG covering 43 plastid genes in eleven wheat genomes were identified (**Figure 5D**). A few photosynthesis and ribosomal protein-related NPGs were retained in the three genomes. For instance, an NPG with a Ks of 0.10 of *petG* was detected in three subgenomes (**Figure S16**), and its ortholog in the wheat A lineage is loss-of-function (**Figure 5E**). A few retained NPGs were lineage specific, such as NPGs of *psbE* in the A lineage, NPGs of *rps8* in the B lineage, and NPGs of *rpl20* in the D lineage. An NPG with a Ks of 0.08 of *psaI* was detected in the wheat A subgenome and *Th. elongatum* genome (**Figure S17**), showing that it may have been integrated into the nucleus before the differentiation of wheat and *Th. elongatum* lineages. Second, we identified a total of 544 retained NMGs covering 24 mitochondrial genes in eleven wheat genomes (**Figure S18A**), and found that a few NMGs were retained in three subgenomes, such as NMGs of *atp9* and NMGs of *nad3*. A few retained lineage-specific NMGs were found, such as NMGs of *atp1* and NMGs of *ccmB* in the D subgenome. An NMG with a Ks of 0.10 of *rpl5* was found only in the B subgenome of three accessions (**Figure S18B-D**).

After integration, most NOGs were pseudogenized during evolution, while a few genes maintained coding abilities to the present. For example, NOGs (**Figure S19A and S19B**) of photosynthesis and translation-related genes tend to maintain coding abilities in wheat over evolutionary time. We also found that the NOGs more likely to maintain coding abilities were generally lineage-specific (**Figure S19C and S19D**). In brief, bread wheat selectively retained more NOGs, which are likely involved in the photosynthesis, translation, and ATP synthase pathways.

### Interactive platform for exploring NOGs in Poaceae

Finally, to explore the dynamic trajectories of NOGs in Poaceae species, we implemented an interactive platform, pNOGmap, for the community to readily access NOG resources (**Figure 6A**). The gene browse tool provides comprehensive information about NPGs and NMGs, including genome location, original organelle gene, NOG cluster information, and collinearity information. We also implemented two interactive microcollinearity tools, including the NPG-MicroCollinearity and NMG-MacroCollinearity tools, for the community to investigate the trajectory dynamics of NPGs and NMGs over evolutionary levels, respectively. Using these two microcollinearity tools, users can track the transfer, elimination, and retention of NOGs in Poaceae and click on a gene to quickly acquire gene information in detail. In total, pNOGmap establishes Poaceae NOG map profiling as a powerful resource for the discovery of functional NOG and their evolutionary history.

**Figure 6.**
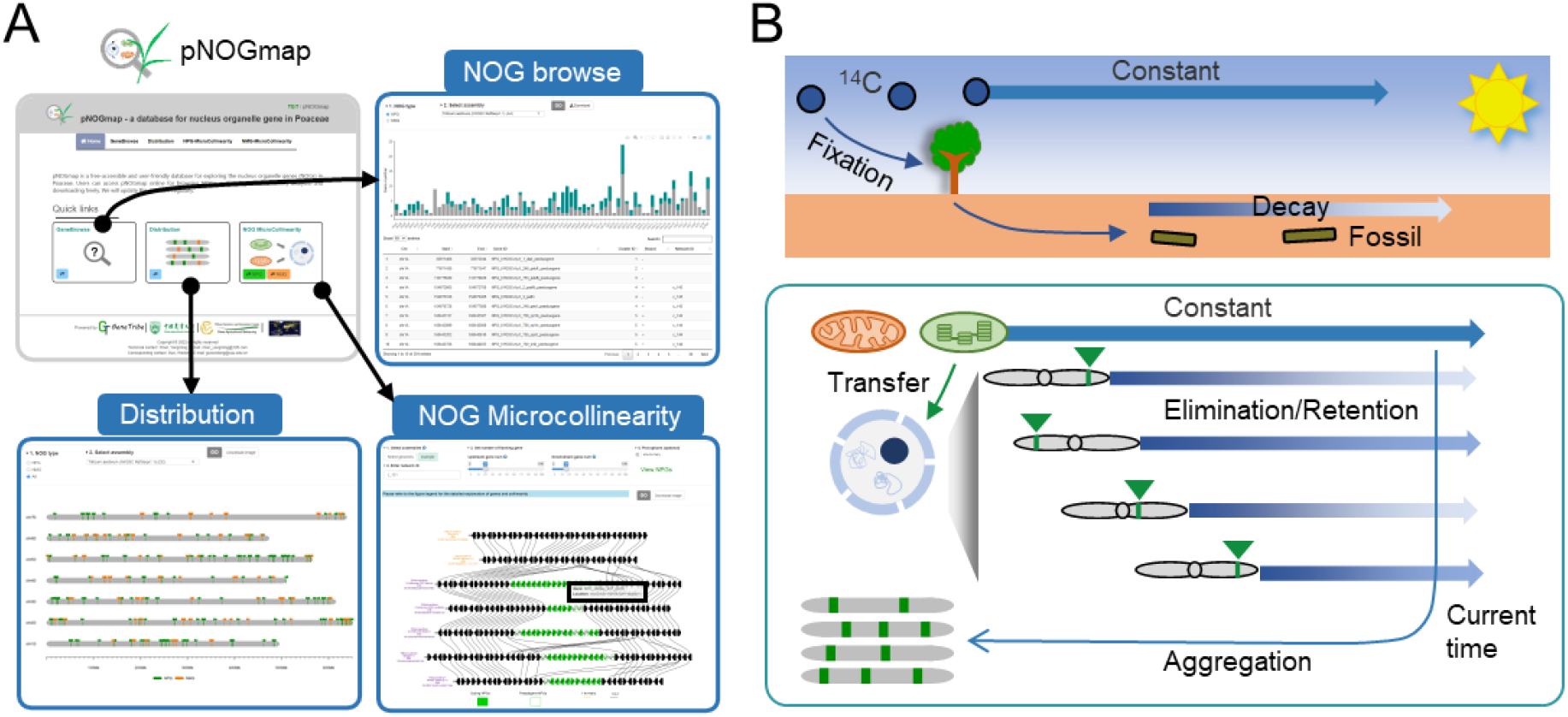
Schematic representation of the Poaceae NOG platform and the radiocarbon-like model of IGTs. (**A**) A snapshot of the pNOGmap database. The gene browse page shows comprehensive information about NOGs, and the distribution page shows the location of NOGs in the genome. Microcollinearity tools show the gene context of NPGC on a local scale across multiple genomes. With this tool, users can trace the origin, elimination, and retention dynamics of NOGs. The detailed gene information is displayed by clicking on the gene (black box). (**B**) A schematic summary showing the radiocarbon-like model illustrating the NOG map recorded the dynamics of organelle-to-nucleus gene transfer and elimination. In plants, organelle-to-nucleus gene transfers ubiquitously and continually occur, while these sequences could undergo elimination during evolution. The process is similar to the decay of carbon-14 in fossils. The observed IGTs at the moment in organisms are aggregated from selective retention and recent transfers over evolutionary time.

## DISCUSSION

A substantial amount of organelle DNA has been continually integrated into the nucleus since the origin of organelles in plants and animals (5, 7). Emerging genome assemblies provide unique opportunities for investigating the roles of IGTs in shaping the inter- and especially intraspecies genome variation and driving plant evolution, but the dynamic trajectories of IGTs at the pangenomic level remain elusive. We developed a method, IGTminer, based on collinearity and gene reannotation to provide a powerful solution for detecting and connecting IGTs in multiple assemblies. The reannotation pipeline in IGTminer can improve the quality of IGT annotation across multiple genomes, and the collinear IGT search pipeline in IGTminer overcomes the inaccuracy of collinear block searches caused by high-similarity IGTs and thus provides great opportunities for investigating the evolutionary trajectories of IGTs.

The trajectory dynamics of IGTs in organisms at the pangenomic level remains to be determined. The integration of IGTs into nucleus genomes introduces PAVs on the local scale and copy number variations on the genome-wide scale, representing a source of intraspecies variation. However, previous studies on IGTs have mainly focused on a single reference genome of a species. Poaceae includes many economically important cereal crops, such as wheat, barley, rye, rice, and maize. We constructed a NOG map in Poaceae using important cereal crops by connecting 67 assemblies. Unlike previous studies (24, 78), the Poaceae NOG map is the first organelle-to-nucleus gene transfer study in plants at the pangenomic level. Extensive collinearity relationships in the Poaceae NOG map were utilized to trace the transfer, elimination, and retention of NOGs over evolutionary time. We found that most IGTs were recently acquired from organellar genomes in the Poaceae. The sequences acquired from organellar genomes were not under the evolutionary constraint seen within the organelle anymore and mutates at a higher rate than the organellar genome. Because the lack of a public platform limits the effective utilization of many genomic resources, we developed pNOGmap, an online database for the community to explore the Poaceae NOG map. In addition, it is a challenge to investigate NOGs if a reference genome or pangenome resource of an interested species is not available or of low-quality. Only four available mitochondrion genomes (wheat, barley, rice, and maize) could be used to infer NMGs, while these four species have pangenome resources and represent diverse genera in Poaceae, and thus can be used to investigate the dynamics of NMGs. More mitochondrial genomes are needed in future studies.

Hexaploid wheat, widely grown worldwide, has undergone two rounds of polyploidization. For Poaceae, we found that both *O. sativa* and *B. distachyon* have more plastid-to-nucleus gene transfers than *A. strigosa*, although the former has much smaller genome sizes. The result is inconsistent with a previous report that a positive correlation existed between nuclear genome size and the total number of NUPTs (10), suggesting that Poaceae species may have different transfer characteristics. We found that large amounts of NOGCs were retained in wheat during two rounds of polyploidization, which can serve as a polyploidization trajectory to support the established evolutionary history of wheat (56, 70). Future studies need more ancestral genomes of wheat to obtain a further understanding of ancestral NOGs. In addition, bread wheat is a staple food crop worldwide compared to durum wheat. Joining of the D genome with more NOGs may contribute to the environmental adaptation of bread wheat, which requires further verification. We also performed pan- and core-NOGC set analysis and revealed the NOG diversity across the wheat population. Private NOGs have lower Ks than core NOGs due to recent transfers. In addition, more selective retention of NOGs in wheat revealed that wheat may need more functional NOGs in an unknown way to maintain growth and development. Organelle genome transformation and editing have been used to advance crop improvement (29, 60). In fact, this study based on pangenome resources shows that organelle-to-nucleus gene transfer continually reshapes the nucleus, which provides more functional copies and genetic diversity for the nucleus. Wheat has more transfers and retention of NOG than maize and rice. This work has connected synthetic biology and genome evolution to some extent.

In plants, organelle-to-nucleus gene transfers ubiquitously and constantly occur and reshape the nucleus, while these sequences tend to undergo elimination during evolution (7, 10). We proposed a radiocarbon-like model illustrating how the NOG map recorded the dynamics of organelle-to-nucleus gene transfer and elimination (**Figure 6B**). Before integration, the genetic material in organelles maintains a low variation rate (79), as carbon-14 is stable in the upper atmosphere. Once integrated into the nuclear genome, organellar-to-nucleus genetic material transfer could undergo elimination and mutation (7, 10), such as the decay of carbon-14 in fossils. Organelle-to-nucleus gene transfer is surprisingly frequent and is a naturally occurring process that pervades nuclear dynamics throughout the evolution of organisms (1, 5). The present NOG map aggregates many nonrandom retentions and recent transfers over evolutionary time. We can also serve the NOGs transferred from different timepoints as the timers characterizing the evolutionary process of species.

Our study developed a robust approach, IGTminer, to identify IGTs and explore their evolutionary dynamics across multiple assemblies based on collinearity and reannotation information. We used IGTminer to generate the first atlas recording the dynamic trajectories of IGTs in plants at the pangenomic level. Our study provides the starting point for investigating IGTs in the pangenome era, expanding our understanding of the roles of IGTs in shaping the inter-and especially intraspecies and driving plant genome evolution.

## METHODS

### Genome resources

We collected 67 sequenced genomes covering 15 species from Poaceae, which represent lineages of both major BEP and PACMAD clades of grasses and cover important crops (**Table S1**). The reference genome sequences (FASTA format) and gene annotations (GFF format) were gathered for each assembly. Assemblies and annotations of *T. aestivum* cv. Zang1817 (41), wheat 10+Genome project (38), *T. urartu* (80), *Ae. tauschii* pangenome (81), *S. cereale* (57, 82), *Th. elongatum* (70), *H. vulgare* pangenome (39), *Av. strigosa* (58), *O. sativa* (40, 60), and *Z. mays* (25, 83) were downloaded from public resources. Raw data for the remaining assemblies were acquired from Ensembl Plants (http://plants.ensembl.org/) and NCBI (https://www.ncbi.nlm.nih.gov/). The genome sequences and annotations were parsed using GffRead (84) and BEDOPS (85).

### Development of the IGTminer workflow

We developed a robust tool, IGTminer, to investigate the dynamics of IGTs at the pangenomic level. The IGTminer workflow was used to incorporate gene reannotation and collinearity information to construct NOG maps. We obtained primary organelle-derived genome sequences as the input of the reannotation pipeline. The genes were classified as pseudogenes according to the following criteria: (1) lack of a short start codon; (2) premature stop codons; and (3) short open reading frame. A polyploid genome was decomposed into multiple diploid subassemblies in advance. We then applied an advanced method to improve the accuracy of collinear NOG search. Multiple adjacent NOGs (NOGC) were regarded as one single integration record and then added to the reference genome. MCScan (86) was used to perform pairwise collinear block detection. The following criteria were used to define homologous NOGCs: 1) at least one shared gene between NOGCs, and 2) these NOGCs are collinear. MCL (53) was used to construct ocollinear networks of NOGCs with an inflation of 2.

### pNOGmap database implementation

The pNOGmap database integrated the Poaceae NOG map and 67 genomes in this study, and implemented multiple versatile analysis and visualization tools for exploring NOGs in Poaceae. It is performed on the Linux operation system and implemented based on the R/shiny framework with a series of R packages, including RMySQL, shinyBS, shinyWidgets, shinycssloaders, DT, plotly, and ggplot2. Homology and annotation datasets were acquired from the TGT database (87). All datasets are deposited in tables managed by a MySQL database engine.

### Divergence time estimation and species phylogeny

The single-copy orthologs from *T. aestivum, T. dicoccoides, Ae. tauschii, T. urartu, Th. elongatum, S. cereale, H. vulgare, As. strigosa, B. distachyon, O. sativa*, and *Z. mays* were identified by GeneTribe (41). A polyploid genome was decomposed into multiple diploid subassemblies in advance. The transcript with the longest coding sequence was used to represent a gene. For each pair of orthologs, their proteins and CDSs were aligned using MUSCLE (88). Then, we converted the protein alignment into a codon alignment using PAL2NAL (89). The Ks of gene pairs was calculated based on the NG model in KaKs_Calculator2.0 (90). The estimation of divergence times between different nuclear genomes was based on the median Ks using a substitution rate of 6.5×10 ^−9^ mutations per site per year. To estimate the transfer times of NOGs, Ks was calculated using the protein and CDSs of NOGs and their original organelle genes. The estimation of the transfer times of NPGs and NMGs was based on the Ks. The rate of substitution of plastids and mitochondria is one-quarter and one-twelfth of that of the nucleus, respectively (79, 91). Phylogenetic trees were inferred based on the PAV matrix of NOGCs using the R package ape.

### Homology inference

A polyploid genome was decomposed into multiple diploid subassemblies in advance. GeneTribe (41) was employed to perform homology inference to generate one RBH (reciprocal best hit) table, two SBH (single-side best hit) tables, two singleton tables, and two ‘1-to-many’ tables. It has been demonstrated that GeneTribe is more accurate than other commonly used homology inference methods among genetically similar genomes. We performed all-vs.-all homology inference for diploid assemblies. Using the same method as Triticeae-GeneTribe (TGT), the microcollinearity plots were visualized in R.

### Detection of NOGs using whole-genome resequencing data

WGS data were acquired from previous studies (41, 75, 76). We used a total of 418 bread wheat accessions, including 170 accessions of landraces, 174 cultivars, and 74 semiwild wheat accessions. The alignment method was described in our previous study (41). Only accessions with reads spanning the junction site between the nuclear genome and NOGC DNA sequence were considered to carry the NOGC.

### Growth conditions and DNA extraction

*T. aestivum* cv. CS and Jagger were used for the experiments. The sterilized seeds were soaked in water for germination at 4 °C for 3 days in the dark, and the seedlings were grown in a growth chamber with growth conditions of 22° C/18 °C (day/night), 16 h/8 h (light/dark), and 50% humidity. The seedlings grew for 4 days in the glasshouse. Approximately 1.5 μg of genomic DNA was isolated from young leaves of both CS and Jagger using the CTAB protocol.

### PCR amplification

To prove that plastid and mitochondrial fragments were indeed inserted into the nuclear genome of CS or Jagger, we designed primers at flanking regions of organelle DNA-nucleus junction sites. The primers were designed using DNAMAN, and are listed in Table S3. A total of four NPGC and NMGC junction products were amplified from the total extracted genomic DNA. The PCR products spanned organelle DNA-nucleus junction sites and were validated by first-generation sequencing.

## AUTHOR CONTRIBUTIONS

W.G., H.P., and Q.S. conceived and supervised this work. Y.C. designed and wrote the program. Y.C., Y.G., X.X., Z.W., L.M., and Z.Y. performed analysis. Y.G. performed experiments. Y.C., Y.J., C.X., J.L., Z.H., M.X., Y.Y., Z.N., Q.S., H.P. and W.G. interpreted data. Y.C. wrote the manuscript, and W.G. and H.P. revised it. All authors read and approved the final manuscript.

## ACKNOWLEDGMENTS

We thank Zhen Qin, Wenxi Wang, and Kuohai Yu for technological support. We thank Longhui Bai and Prof. Handong Su for helpful discussions and technological support.

